## Supplementary Figures for "Drosophila SWR1 and NuA4 complexes are defined by DOMINO isoforms"

### Figure S1

## A

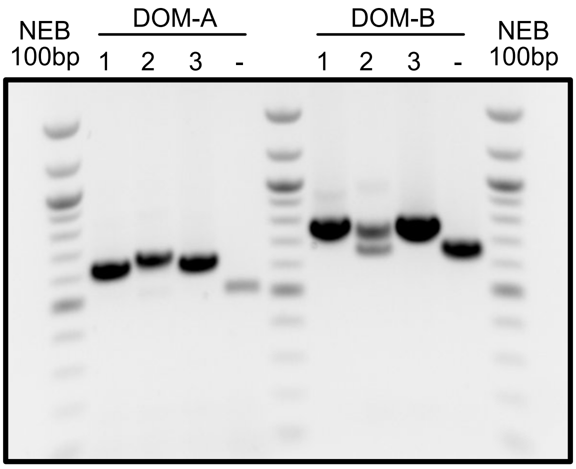

### Figure S2

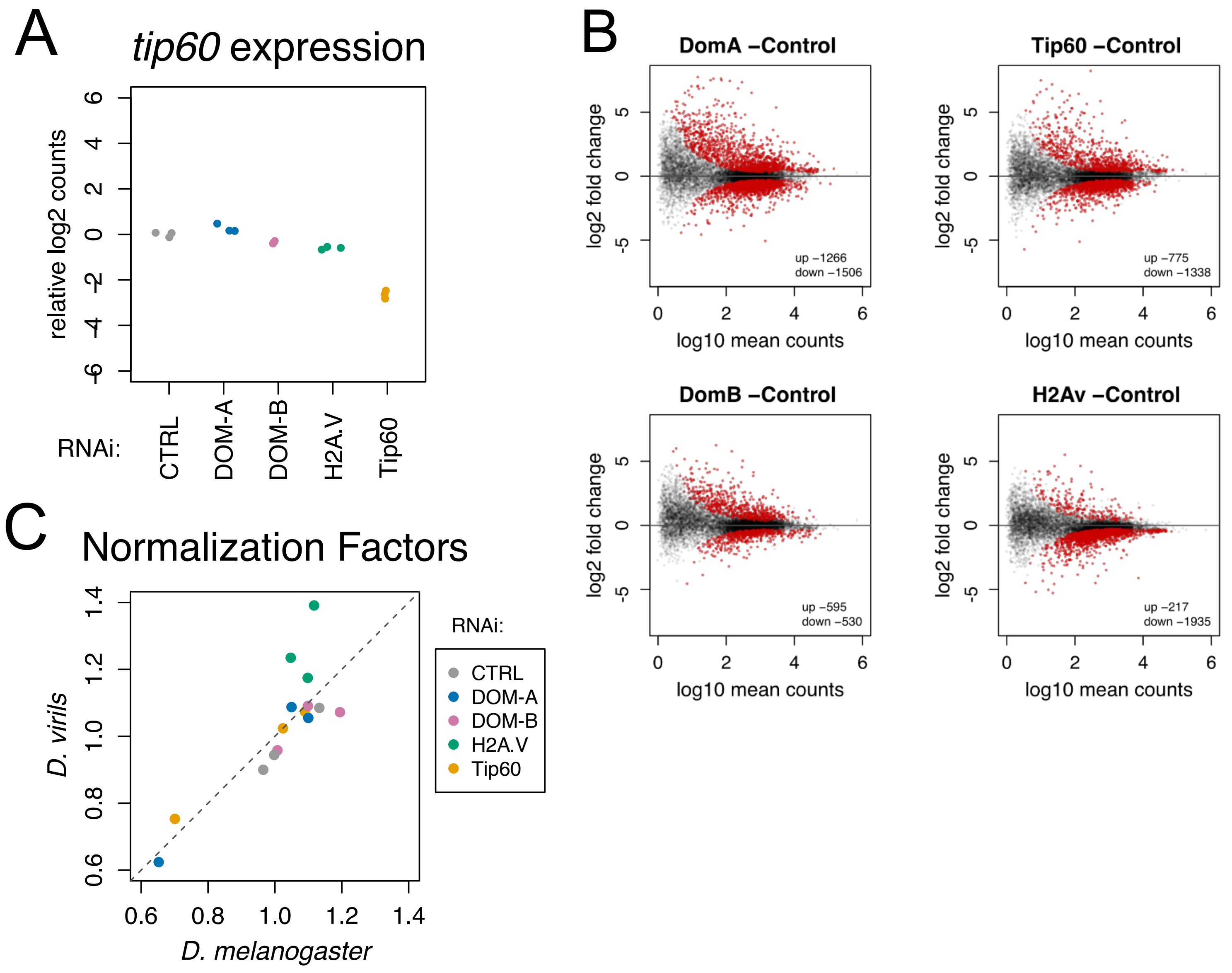
A
tip60 expression
relative log2 counts
6
4
2
0
-2
-4
-6
RNAi:
CTRL
DOM-A
DOM-B
H2A.V
Tip60
B
DomA –Control
Tip60 –Control
DomB –Control
H2Av –Control
log2 fold change
5
0
-5
log10 mean counts
0
2
4
6
up –1266
down –1506
up –775
down –1338
up –595
down –530
up –217
down –1935
C
Normalization Factors
D. virils
D. melanogaster
0.6
0.8
1.0
1.2
1.4
RNAi:
CTRL
DOM-A
DOM-B
H2A.V
Tip60

### Figure S3

A

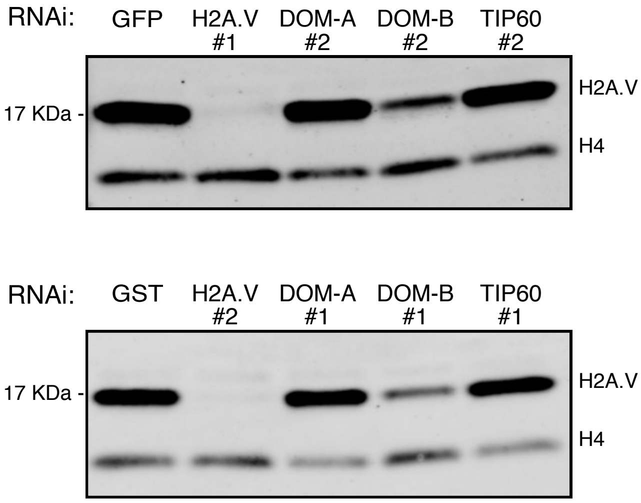


B

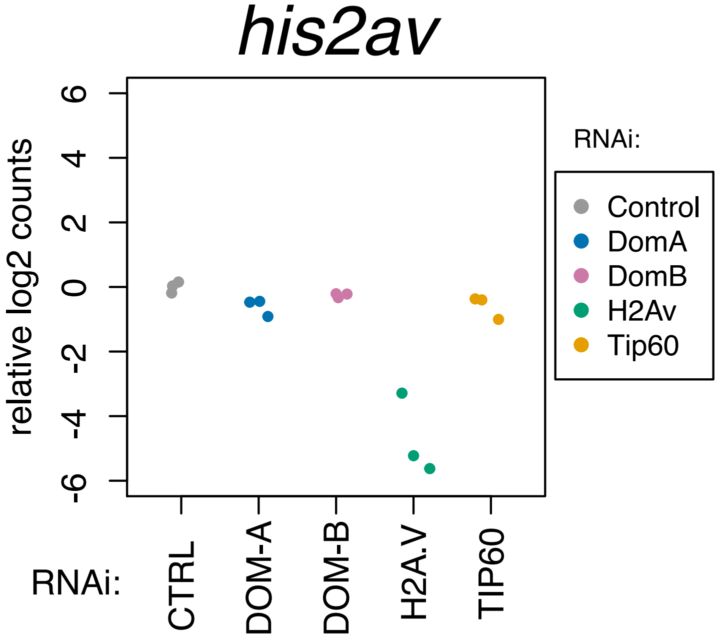


C

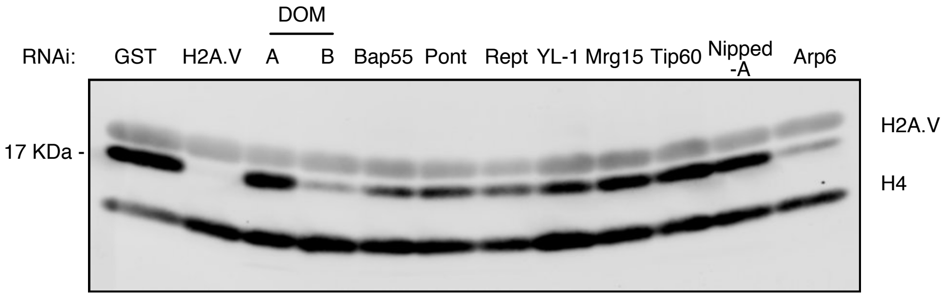


D

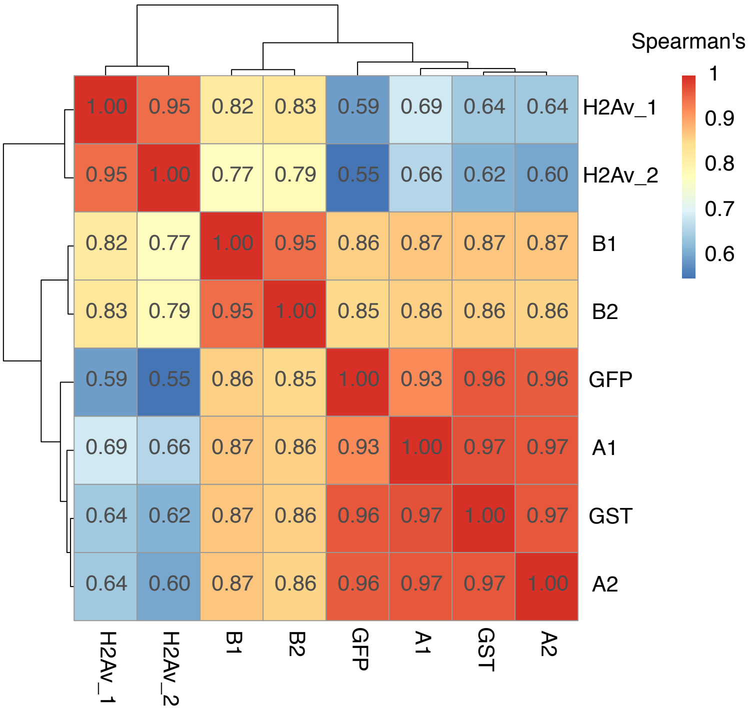


E

| Comparison | p-value |
|---|---|
| DOM-B_WT_DOM-B vs DOM-B_WT_CTRL | 0.000102860598845281 |
| DOM-B_KG_DOM-B vs DOM-B_KG_CTRL | 0.094182124842038 |
| (DOM-B_WT_DOM-B vs DOM-B_WT_CTRL) vs (DOM-B_KG_DOM-B vs DOM-B_KG_CTRL) | 0.00248791036943397 |

### Figure S4

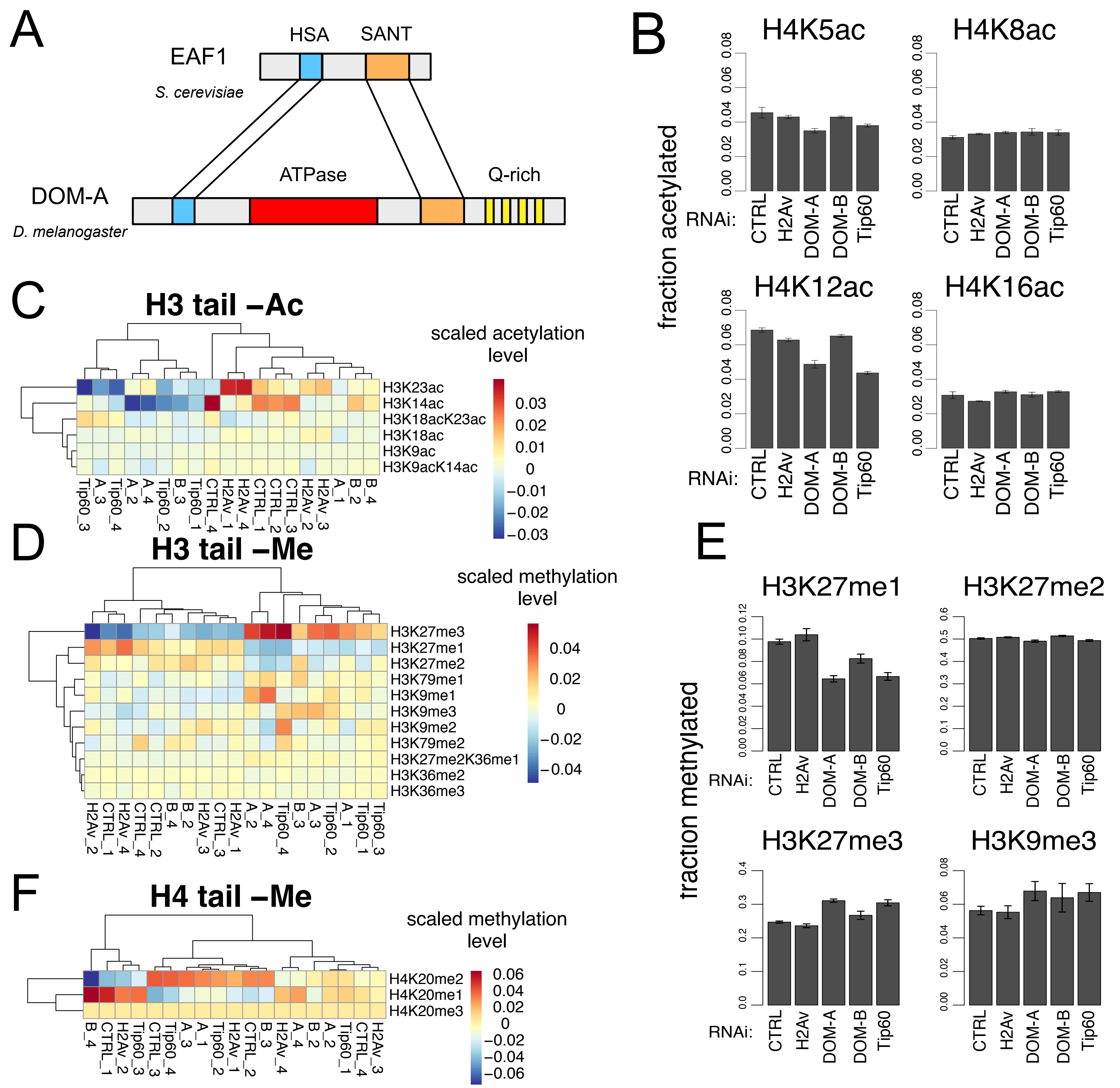


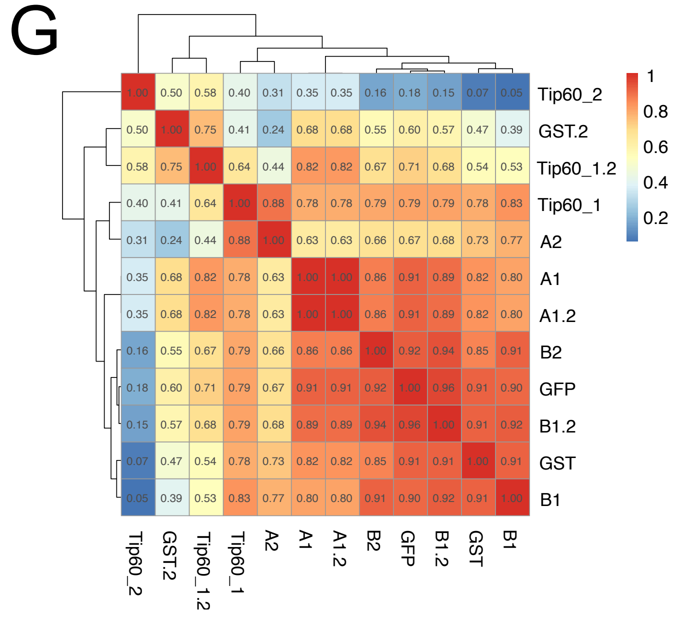


H

| Comparison | p-value |
|---|---|
| DOM-A_WT_DOM-A vs DOM-A_WT_CTRL | 0.00376506093514611 |
| DOM-A_KG_DOM-A vs DOM-A_KG_CTRL | 0.00101987027757499 |
| (DOM-A_WT_DOM-A vs DOM-A_WT_CTRL) vs (DOM-A_KG_DOM-A vs DOM-A_KG_CTRL) | 0.374967301291081 |
