## Supplementary Figure Legends for "Drosophila SWR1 and NuA4 complexes are defined by DOMINO isoforms"

**Supplementary Figure 1**

A. Agarose gel analysis of CRISPR clones. The insertion of a 3XFLAG tag results in a PCR product 72bp longer than in the untagged cells (CTRL). The presence of two bands indicates heterozygosity. NEB 100 bp ladder servers a size marker.

**Supplementary Figure 2**

A. Plot showing relative log2-expression of Tip60 mRNA as measured by RNAseq upon knockdown of protein indicated on the x-axis and by color-coding. The values of 3 biological replicates are shown in the plot.
B. Differential gene expression analysis on coding genes from RNA-seq data. Scatter plots represent log_2_ fold-change of knock-down indicated in the plot title over CTRL knock-down for each gene (N=10250) in relation to its mean expression (log10 mean counts). Red dots represent significant (adjusted p-value <0.01) up- or down- regulated genes.
C. Scatter plot comparing normalization factors used to scale transcriptomes calculated either from the *D. melanogaster* or the *D. virils* (spike-in) genome. A deviation from the diagonal line indicates global upregulation (lower *D. virils* scale factor) or downregulation (higher *D. virils* scale factor) of transcription. The values of the 3 biological replicates are shown in the plot. Samples from the different knock-down samples are color-coded.

**Supplementary Figure 3**

A. Replicate western blot showing the expression of H2A.V in nuclear fractions derived from cells treated with dsRNA against GST (CTRL), H2A.V (H2A.V), DOM-A (A), DOM-B (B) and Tip60. Histone H4 (H4): loading control.
B. Western blot showing the expression of H2A.V in nuclear fractions derived from cells treated with dsRNA against some of the DOM interactors identified by mass-spectrometry (common between DOM-A and DOM-B or specific for one or the other isoform). Histone H4 (H4) serves as loading control.
C. Heatmap showing Spearman’s correlation coefficients between replicates and dsRNA treatments calculated for H2A.V ChIPseq normalized coverage signal around the Transcription Start Site (TSS) (± 2000 bp). Clustering is based on Euclidean distance. Individual biological replicates are shown.
D. Table listing calculated p-values (linear regression) for the comparisons depicted in Figure 3G. Naming structure is the following: transgene (WT or KG) RNAi

**Supplementary Figure 4**

A. Schematic comparison of S. cerevisiae Eaf1 and D. melanogaster DOM-A. Conserved domain structure is highlighted,
B. Barplot showing the average fraction of acetylated peptide (over non-acetylated) (n=3 for B, n=4 for all the others) for 4 residues in histone H4 tail (K5, K8, K12, K16) upon knock-down of the proteins indicated on the x-axis. Error bars represent SEM.
C. Heatmap shows scaled acetylation levels for various histone H3 residues (measured by mass-spectrometry) in cells treated with dsRNA against GST or GFP (CTRL), H2A.V (H2A.V), DOM-A (A), DOM-B (B) and Tip60. Individual biological replicates are shown. Rows and columns are clustered based on Euclidean distance.
D. Same as (c.) but showing scaled methylation levels for various histone H3 residues
E. Barplot showing the average fraction of methylated peptide (over non-acetylated) (n=3 for B, n=4 for all the others) for H3K27 upon knock-down of the proteins indicated on the x-axis. Error bars represent SEM.
F. Same as (c.) but showing scaled methylation levels for histone H4K20 residue
G. Heatmap showing spearman’s correlation coefficients between replicates and dsRNA treatments calculated for H4K12ac ChIPseq normalized coverage signal around the Transcription Start Site (TSS) (± 2000 bp). Clustering is based on Euclidean distance. Individual biological replicates are shown.
H. Table listing calculated p-values (linear regression) for the comparisons depicted in Figure 4E. Naming structure is the following: transgene (WT or KG)_RNAi

**Supplementary Table 1**

Excel spreadsheet containing LFQ values obtained from the MaxLFQ algorithm and *limma* output.

**Supplementary Table 2**

Result tables from DEseq2 analysis.

**Supplementary Table 3**

gRNAs, repair templates and primers used in this study.
